# Long-Term Expression and Safety of AAV1-Mediated PI3Kδ Overexpression in the Adult Rat Cortex

**DOI:** 10.64898/2026.01.22.700520

**Authors:** Lydia Knight, Zuzana Polčanová, Dana Mareková, Lucia Machová Urdzíková, Daniel Jirák, Natalia Ziolkowska, Pavol Makovický, Fred de Winter, Jessica Kwok, Pavla Jendelová, James Fawcett, Kristýna Kárová

## Abstract

Mature CNS neurons are incapable of sufficiently regenerating their axons following spinal cord injury (SCI). This is largely due to developmental changes in epigenetic control leading to suppression of axon growth transcriptomic profile, leading to a shift towards synapse function support. Recently, manipulating the PI3K/Akt/mTOR pathway through PI3Kδ overexpression in cortical neurons enhanced axonal regeneration of corticospinal tract axons, which was accompanied by functional recovery monitored for up to 16 weeks. However, PI3K is more widely known for its role as an oncogene, and since overexpression is achieved by the use of AAVs, valid safety concerns are raised as it is unknown what the long-term consequences of sustained PI3Kδ expression in the brain are, which may be necessary to achieve complete re-establishment of the motor pathway. In this study, AAV1-hSYN-PIK3CD was injected into the motor cortex of rats, which survived for 1 year. Comparison with uninjected control animals reveal stable PI3Kδ expression and sustained pathway activation through increased pS6. PI3Kδ-treated animals show absence of tumour formation, neural soma hypertrophy, glial cell activation, or haematological or biochemical abnormalities. Thus, long-term neuronal PI3Kδ expression appears to be well tolerated and may provide a safe and durable strategy to promote functional repair following SCI.

## Introduction

Injuries to the adult central nervous system (CNS) frequently lead to permanent functional impairments, largely attributed to the marked decline in the intrinsic regenerative capacity of mature neurons, accompanied by an extrinsically remodelled inhibitory environment that suppresses regeneration. These remain major barriers to recovery as mature neurons are incapable of long-distance axonal growth and therefore are unable to re-establish functional connections. The decline in intrinsic regenerative capacity is due to a multitude of factors that occur in both the soma and the axon. For example, a gradual decline in the expression of receptors (such as integrins) associated with growth and migration (Stepankova et al, 2025). Another key molecular contributor to this intrinsic decline is the developmental downregulation of class I phosphatidylinositol-3-kinases (PI3K), specifically the δ isoform. PI3Ks are activated by a variety of ligands such as growth factors binding to transmembrane receptor tyrosine kinases (Algeria et al, 2018). PI3K’s are a family of enzymes that catalyse the phosphorylation of phosphatidylinositol (4,5)-biphosphate (PI (4,5) P_2_) at the 3’ position of the inositol ring to generate the signalling lipid phosphatidylinositol (3,4,5)-triphosphate (PIP_3_) (Anderson and Jackson, 2003). PIP_3_ serves as a membrane-localised docking site for downstream effectors containing a plekstrin homology (PH) domain, which activate downstream signalling pathways such as the Akt/mTOR. This pathway is known to govern diverse cellular processes including growth, survival, and metabolism (Carnero *et al*, 2008; Manning and Cantley, 2007; Glaviano et al, 2023). Additionally, PI3Kδ and its various isoforms play a fundamental role in maintenance of both adaptive and innate immunity through B and T cell signalling, monocyte survival and differentiation, erythropoiesis, cytokine production, and platelet activation (So and Fruman, 2012; Tothova et al, 2021, Lanahan et al, 2023). A gain of function mutation in the PIK3CD gene encoding the p110δ catalytic subunit results in an immunodeficiency disease called activated phosphoinositide 3-kinase δ syndrome (APDS). Other mutations can also lead to increased PI3Kδ enzyme potency and may result in effector T cell senescence and lymphoproliferation (Singh et al, 2020).

Manipulating the PI3K pathway to promote CNS axonal regeneration gained recognition in 2009, when genetic deletion of PTEN in mice with optic nerve injury led to robust axon regeneration (Park *et al*, 2009). PTEN is a lipid-phosphatase that reverses the PIP_2_ to PIP_3_ transition and so opposes the action of PI3K (Carracedo and Pandolfi, 2008). Blocking PTEN function therefore leads to increased PIP3 levels and mTOR pathway activation. It has more recently been shown that PIP_3_ levels are high during CNS development and its decline with maturity corresponds to the loss of CNS regenerative capacity (Nieuwenhuis *et al*, 2020). This PIP3 decline in mature neurons, likely due to reduced expression of PI3K isoforms, makes PTEN knockouts less effective in adults (Geoffroy *et al*, 2016). Although high titre AAV driven overexpression of constitutively active Akt3 in the CST successfully upregulated mTOR signalling and induced robust regeneration, this was also accompanied by behavioural seizures, as well as neuronal soma hypertrophy and hemimegalencephaly, confounding functional outcomes (Campion et al, 2022). Notably, excessive or dysregulated activation of the PI3K-Akt-mTOR pathway has been strongly implicated in epilepsy and cortical malformations, with sustained mTOR signalling driving abnormal neuronal growth, altered network excitability, and seizure susceptibility in both experimental models and human neurodevelopmental disorders (Crino, 2015; Limanaqi et al, 2020; Lasarge and Danzer, 2014).

We recently conducted a regeneration study in adult rats that received C4 dorsal column crush spinal cord injury (SCI) with AAV1-mediated delivery of naturally hyperactive PI3Kδ to their right sensorimotor cortex, with the aim to regenerate axons of the injured corticospinal tract (CST). Behavioural tests showed significant improvements in treated rats, accompanied by regrowing axons up to 1cm below the lesion site and without neuronal hypertrophy or seizures (Karova *et al*, 2025).

Growing evidence supports a promising role for PI3K in promoting CNS regeneration, leading to increased interest in therapeutic strategies that involve PI3K overexpression or activation. However, it is critical to assess the long-term safety and potential tumorigenic consequences of sustained PI3K activation in the brain cortex. One of the downstream effects of PI3K pathway activation is cell survival and proliferation, through downstream AKT phosphorylation which inactivates the pro-apoptotic transcription factors FOXO and BAD. Further PI3K/Akt/mTOR pathway effectors, ribosomal protein S6 kinases (S6K) and eukaryotic translation factor 4E binding protein (4E-BP1) are phosphorylated, allowing the initiation of protein synthesis (He et al, 2021; Manning and Toker, 2018; Nelson et al, 2024). It is well established that PI3K is an oncogene, and to date, 5 PI3K inhibitors have already been approved by the FDA for cancer treatment (Trigueiros et al, 2024). Gain of function mutations in the gene encoding PI3Kδ (PIK3CD) drives haematological cancers through sustained malignant B-cell proliferation and resistance to apoptosis, as in patients with indolent non-Hodgkin’s lymphomas (Gopal *et al*, 2014). Moreover, Akt activation can directly disrupt cell-cycle checkpoints by phosphorylating and misplacing the CDK inhibitor p27^Kip1^, preventing its nuclear inhibition of cyclin– CDK complexes and thereby promoting uncontrolled proliferation (Shin et al., 2002). Aberrant activation of the PI3K signalling cascade is observed in ∼88% of glioblastomas (Langhans *et al*, 2017). Therefore, to reduce the potential of PI3K induced oncogenic effects, we deploy the neuronal specific hSYN promoter. However, valid safety concerns need to be addressed to confirm that long-term expression of PI3Kδ does not lead to tumour formation.

The use of adeno-associated virus (AAV) for experimental CNS gene therapy has become the standard route of delivery with one AAV type 2 mediated gene therapy applied in the brain putamen receiving accelerated approval by the FDA for the treatment of AADC deficiency (ClinicalTrials.gov Identifier: NCT02852213, 2025). In the context of cortical and corticospinal motor neuron (CSMN) transduction, a comparison of four different AAV serotypes revealed that AAV1 is optimal following rat sensorimotor cortex injections (Hutson et al, 2011), favouring its use in studies since. Previous, though limited, studies have demonstrated that AAV1 can drive long-term expression in the rodent CNS. For example, intraventricular AAV1-HβH injection in neonatal mice resulted in detectable enzyme expression in the brain one year later (Passini et al, 2003). However, the majority of studies to date have evaluated much shorter durations, typically only extending to a few weeks or months (Karova et al, 2025; Stepankova et al, 2025; Blits et al, 2003; Chen et al, 2007; Huang et al; 2011). AAVs have a good reputation in terms of safety, particularly when used with a neuron-specific promoter such as hSYN where off-target effects are considered mild compared to other gene delivery methods (Au et al, 2022). Off-target and unwanted effects such as innate and adaptive immune activation, vector neutralisation or rare vector integration events have been reported, particularly at high vector titres (Selot et al, 2014; Reichel et al, 2017; Rogers et al, 2011; Batty and Lillicrap, 2024).

In this study, rats received intracortical injections of AAV1-hSYN-PI3Kδ and survived for one year without SCI, to address the following questions: 1.) Given that AAV1-hSYN-PI3Kδ drives robust regeneration and strong transgene expression at 16 weeks, would the same recombinant vector express as strongly after one year? 2.) Does long term AAV1-hSYN-PI3Kδ expression result in any adverse local or systemic effects? Addressing these questions provide important insight into both the stability of AAV1-hSYN-mediated expression and the safety of sustained neuronal PI3Kδ overexpression.

## Methods

### Animals

All experiments were performed in accordance with the European Communities council directive of September 22, 2010 (2010/63/EU), following the ARRIVE guidelines (https://arriveguidelines.org/), and were approved by the Ethics Committee of the Institute of Experimental Medicine ASCR, Prague, Czech Republic.

Healthy 10-week-old Wistar rats were used in this study. Both male and female rats were used to cover potential sex differences. Throughout the study, rats were housed in pairs with a 12h light/dark cycle and standard conditions, provided with water and food *ad libitum*. They were quarantined for 2 weeks prior to intracortical injections, after which they were checked every day with increased care taken for the first 3–5 days. Rats were routinely assessed for overall health status, with their body weight measured on a weekly basis. Animals were then sacrificed after 1 year.

### Viral vectors preparation

Plasmid DNA encoding AAV-SYN-PIK3CD (Addgene plasmid #203730) was made by VectorBuilder. Plasmid DNA was scaled in house using DH5α transformation competent cells (Invitrogen, Waltham, MA, USA) and isolated with Maxiprep (ThermoFisher, Waltham, MA, USA). Subsequently, correct ITR sites were verified with a restriction analysis and electrophoresis. Plasmids were then used to generate AAV1-hSYN-PIK3CD, titre 2.7×10^12^gc/mL according to a previously described protocol (Verhaagen et al, 2017). AAV1-SYN-eGFP was purchased from Vigene Biosciences with a titre of 2.3 × 10^13^ (now part of Charles River, cat. #CV17001-AV1). Before use, AAV1-hSYN-eGFP was titre matched with AAV1-hSYN-PIK3CD.

### Virus injections

Rats weighing 200-300 g were anesthetised with 3% isoflurane, placed on a heating pad maintained at 37°C, and their body temperature monitored via rectal thermometer. Once anesthetised, they received subcutaneous buprenorphine (Bupaq Multidose 0.3 mg/ml, Richter Pharma, Austria, 0.01 mg/kg) and intramuscular caprofen (Rymadil, Pfizer, 7.5 mg/lg) for analgesia. Animals were secured in a semiautonomous stereotaxic frame (Neurostar, Tubingen, Germany). The scull was exposed, and drill and syringe positions were calibrated relative to bregma and lambda, with adjustments made for tilt and scale as needed. For the experimental group, only one hemisphere was injected, while the contralateral hemisphere served as an internal control. Control rats underwent no surgical procedures or injections. Virus was delivered at four coordinates per injected hemisphere: ML/AP: 1’5, 1;2, 3’5; 3’5, 2;3, 0’5. At each site, 0.5 μL of either AAV1-hSYN-eGFP or AAV1-hSYN-PIK3CD virus was injected at a depth of 1.5mm using a 10μL Hamilton syringe, at a rate of 0.2μL/min. After injection, the needle remained in place for 3 minutes before being retracted in 0.5 mm increments every 30 seconds. Finally, the scalp was sutured and treated with Novikov solution. Each rat received 1 ml of glucose solution intraperitoneally (Ardeanutrisol, Glucose 100 g/l, Ardeapharma, Sevetin, Czech Republic) at the end of surgery.

### Perfusion and tissue processing

At the end of the 1-year survival period, animals were deeply anesthetised by isoflurane and were injected with pentobarbital (50 mg/kg, i.p.). Rats were transcardially perfused with phosphate-buffered saline (PBS) followed by 4% paraformaldehyde (PFA) in PBS. Brains were then removed from the skull and post-fixed in 4% PFA overnight.

### Immunohistochemistry

Brains were incubated in sucrose solution with gradually increasing concentration (10%, 20% and 30% sucrose in deionised water). Tissue was then embedded in OCT mounting media (VWR, #03820168) and sections cut at 40 μm on a cryostat (Thermo Scientific, Cryostar NX70). Floating frozen coronal brain sections were allowed to come to room temperature, then washed 3 times for 10 min each in 1x TBS to remove cryoprotectant. When staining for PI3Kδ and pS6, heat induced antigen retrieval (HIER) was performed prior to the normal staining protocol. Sections were washed once in dH_2_O, then transferred to an aqueous 4.5mM citraconic anhydride (125318, Sigma Aldrich) solution (pH 7.4), which was performed in a fume hood. The plate was placed in a 97 °C water bath and left for 20 minutes. Sections were then allowed to cool completely before washing once more in dH_2_O, 1x with TBS, and continuing with usual protocol. Sections were then permeabilised using 2% Triton in TBS for 20 min. In samples where biotin-streptavidin amplification was used, blocking endogenous avidin/biotin was done using Avidin/Biotin block (Abcam, Bristol, UK, ab64212). Subsequently, sections were blocked for 2h at room temperature using 10% Chemiblocker (Merk Millipore, Billerica, MA, USA, #2170), 0.3M glycine, 0.2% triton, and TBS. Primary antibodies were incubated for 2 days at 4^0^C, in the dark. Samples were then washed and incubated in their respective secondary antibody (1:400, Life technologies, Carlsbad, CA, USA) or biotinylated secondary antibody (1:400, Vector biotechnologies) for 2 hours at 4^0^C, then washed and incubated for an additional 2 hours, at 4^0^C, with Streptavidin Alexa Fluor 488/594 (1:400, Life technologies, Carlsbad, CA, USA) and finally DAPI (1:5000) for 10 min. After washing, sections were mounted with Vectashield fluorsafe (Vector laboratories, CA, USA). Primary antibodies used in this study were rabbit anti-p110 (1:400, Proteintech, Manchester, UK, 21708-1-AP), mouse anti-pS6 (1:400, Cell Signaling, Danvers, MA, USA #62016), chicken anti-GFP (1:400, Thermo Fisher Scientific, A10262), mouse anti-Iba-1 (1:400, Invitrogen, Carlsbad, CA, USA, #GT10312), mouse anti-GFAP-Cy3 (1:400, Merk Millipore, Billerica, MA, USA, #C9205).

For the initial PI3Kδ staining, all brains were processed and stained (n = 10 for uninjected controls, n = 10 for PI3Kδ injected animals). For subsequent staining’s, with H&E, Iba1, GFAP and pS6, 5 animals per group were used. This ensured that sufficient brain sections remained available, since overexpressed PI3Kδ is detectable in limited number of coronal brain sections.

### PI3Kδ, pS6 expression level analysis and soma size

Comparison of mean fluorescent intensity (MFI) of PI3Kδ between uninjected 1-year old control rats (*n* = 5) and rats expressing PI3Kδ for 1 year (*n* = 5), or 16 weeks (*n =* 5) was visualised using 40μm brain coronal sections. Images were obtained under the same settings using a confocal microscope at 20x magnification (Olympus, SpinSR10). Using image J, the PI3Kδ+ region, or equivalent, was selected and MFI analysed. The same was done for sections stained for pS6, (*n =* 5 uninjected controls, *n* = 3 GFP controls, *n =* 5 PI3Kδ injected), and MFI normalised to background. One-way ANOVA with Tukey’s post hoc test or unpaired t-test was used in statistical analysis.

Neuronal soma size was determined using the same images. In ImageJ, the diameter of layer V PI3Kδ or GFP positive neurons was measured using the measure length tool and drawing a straight line across the vertical diameter of the cell. At least 60 neurons were analysed across 3 images per animal (*n =* 5).

### Microglial and astrocyte morphology and microglia cell count

Microglia and astrocyte morphology were analysed using 40 μm coronal brain sections stained for Iba-1 or GFAP (*n =* 5). Images were obtained under the same settings for MFI, at 20x magnification, and optimal settings for Sholl analysis, 60x magnification using a confocal microscope (Olympus, SpinSR10). 7 astrocytes or microglia per image were randomly selected, and 3 images per animal analysed. Branching analysis was done using Scholl analysis in Image J utilising the neuroanatomy plugin. Concentric circles (Sholl shells) placed 3 or 5μm apart measured the number of intersections from the soma. Unpaired t-test or two-way ANOVA with Sidak’s post hoc test was used in statistical analysis.

The same images were used to determine microglial cortical densities. ROIs measuring 500 μm x 500 μm were drawn in three distinct areas of the motor cortex, and Iba1^+^ cells within each ROI were counted using Imaris 10.2. Cell counts are an average of 3 ROIs per image, 3 images per animal, *n =* 5.

### MRI acquisition and analysis

Magnetic resonance imaging (^1^H MRI) was performed to assess brain morphology and detect potential structural abnormalities. Scans were acquired using a RARE (Rapid Acquisition with Relaxation Enhancement) sequence with a repetition time (TR) of 3300 ms. Coronal images were obtained with an averaging (AV) of 2, resulting in an acquisition time of 2 minutes and 38 seconds, a slice thickness of 0.8 mm, covering 15 slices, and a field of view (FOV) of 4.5 × 4.5 cm. Axial images were acquired with an AV of 4, an acquisition time of 5 minutes and 16 seconds, a slice thickness of 0.8 mm, covering 25 slices, and an FOV of 4 × 4 cm. Images were qualitatively assessed. 8 uninjected controls and 10 PI3Kδ animals were assessed.

### Tissue collection and histological analysis

In a terminal setting, liver, spleen, kidneys and lungs were harvested and weighed using a precision weight scale before being further processed for histological analysis. Organs were fixed in 4% paraformaldehyde in 0.1M PBS for 24 hours. Fixed specimens were then embedded in paraffin (FFPE) and sectioned into 5μm slices before being stained with haematoxylin-eosin (Carl Roth, Karlsruhe, Germany, X883.2, CN04-1) following standard protocols to visualise tissue morphology and cellular architecture. Briefly, sections were deparaffinised in xylene, rehydrated through graded alcohols, stained with haematoxylin, differentiated and blued, followed by eosin counterstaining, dehydration, clearing, and mounting.

### Haematology and Biochemistry

At the end of the 12-month experimental period, blood samples were collected. Animals were anesthetized using an inhaled mixture of isoflurane (3% v/v; AbbVie, Chicago, IL, USA) delivered in air at a flow rate of 300 mL/min. Anaesthesia was maintained throughout the duration of blood collection. Blood samples were obtained from the retro-orbital venous sinus by gently inserting a sterilized Pasteur pipette tip into the medial canthus, beneath the nictitating membrane. The pipette was carefully positioned between the eyeball and the bony orbit to facilitate sample collection. Blood samples were collected into BD Microtainer® collection tubes. For haematological analysis, samples were drawn into tubes containing K_2_EDTA (Becton, Dickinson and Company #365974) to prevent coagulation. Haematological markers analysed were: leukocytes, lymphocytes, monocytes, neutrophils, eosinophils, basophils, erythrocytes, red cell distribution width-coefficient of variation (RDW-CV), mean corpuscular volume (MCV), reticulocytes, haemoglobin, mean corpuscular haemoglobin (MCH), haematocrit, and thrombocytes. Blood samples for biochemical analysis were collected in a serum-separating tube (Becton, Dickinson and Company #365963) and the following serum parameters measured: calcium, potassium, sodium, iron, phosphorus, total protein, albumin, and glucose. Biochemistry and haematology results were evaluated by the Czech branch of Synlab (Munich, Germany). Unpaired t-test was used for statistics.

While the animal remained under anaesthesia, urine was collected by gently applying manual pressure to the lower abdomen while holding the rat over a Petri dish. The urine was collected with a syringe into an Eppendorf tube (1.5 mL). Proteinuria, pH, leukocytes, specific density and glycosuria levels were assessed. Similarly, urine analysis was performed by Synlab (Munich, Germany).

## Results

### Stable PI3Kδ expression maintains pathway activation without neuronal hypertrophy

We previously demonstrated that AAV-mediated expression of hyperactive PI3Kδ promotes corticospinal tract (CST) regeneration at 16 weeks post-delivery, with elevated PI3Kδ expression sustained at that timepoint. To evaluate long-term expression of PI3Kδ in the absence of injury, we examined projection-layer cortical neurons in rats with intracortical injections of AAV1-hSYN-PIK3CD into the right motor cortex. At one-year post AAV1-hSYN-PIK3CD injection, immunostaining for PI3Kδ revealed persistent expression in the cortex, including layer V cortical neurons which exhibited significantly elevated mean fluorescent intensity (MFI) compared to controls (P < 0.0001; Figure 1A, C, D). When compared with 16-week expression levels, MFI showed a slight but non-significant decline over time (P = 0.6070; Figure 1B, D).

**Figure 1.**
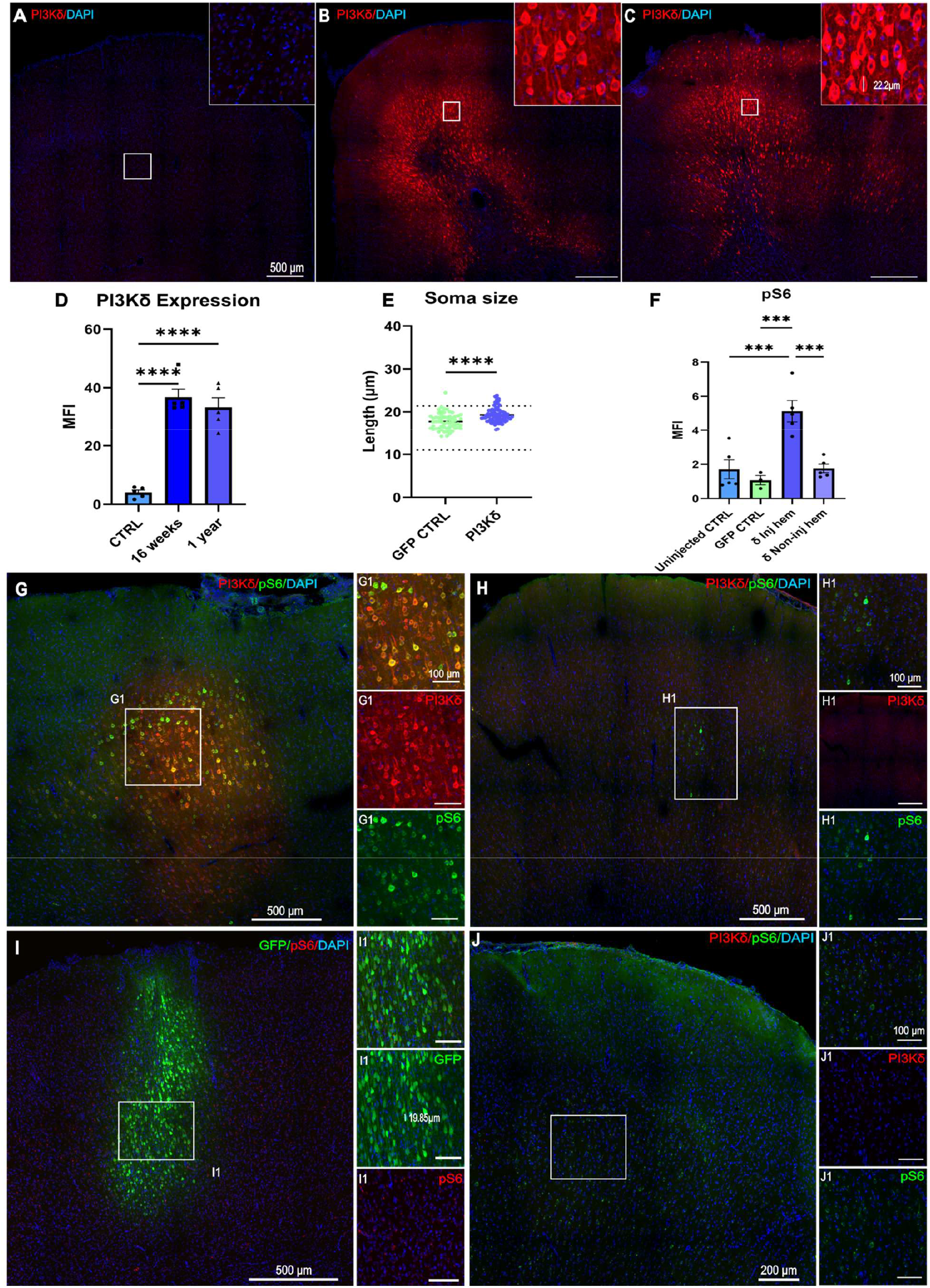
PI3Kδ expression is sustained in injected cortex after 1 year, with elevated pS6 and normal soma morphology. Endogenous PI3Kδ expression in layer V cortical neurons is low in uninjected controls (**A**), and significantly higher in rats injected with AAV1-hSYN-PIK3CD both 16-weeks (**B**) and 1-year after injections (**C**). Quantification shows MFI is similar at both time points (**D**). Lengths of PI3Kδ or GFP-expressing neurons were measured and remain in normal soma size range, although GFP-expressing neurons are significantly smaller (**E**). Phosphorylation of S6 remains upregulated in PI3Kδ-expressing neurons 1 year after injections, quantified by MFI (**F, G**). Significantly lower levels of pS6 were detected in noninjected cortex (**H**) GFP controls (**I**) and uninjected controls (**J**). Magnification 20x. Data are presented as mean ± SEM, ^***^ *p* < 0.001, ^****^ *p* < 0.0001, by T-test or one-way ANOVA with Tukey’s post hoc test. *n* = 5 for uninjected controls, 16 weeks and 1 year PI3Kδ animals, *n* = 3 for GFP controls.

An earlier study using high titre AAV driven AKT3 overexpression to activate the pAkt/mTOR pathway successfully promoted axonal regeneration but also resulted in pronounced neuronal soma hypertrophy accompanied by seizure activity (Campion *et al*, 2022). In contrast, we have previously shown that PI3Kδ treatment does not increase neural soma size above normal level and can partially mitigate SCI-induced reduction in soma size after 16 weeks (Karova *et al*, 2025) highlighting PI3Kδ neuroprotective effect. To assess whether long-term PI3Kδ expression alters soma size in the absence of injury, we measured the soma length and diameter of PI3Kδ^+^ or GFP^+^ layer V cortical neurons one year after brain injections (Figure 1C). The soma measurements in PI3Kδ-expressing neurons were significantly higher than the GFP controls, with mean soma length 19.29 μm and 17.74 μm, respectively (Figure 1C, I, E). However, both remained within the normal size range reported for adult rats as well as previously reported lengths in SCI setting, indicated by the horizontal dotted line (Figure 1E) (Gao and Zheng, 2004, Karova *et al*, 2025).

Phosphorylated S6 (pS6), a downstream target of mTORC1, is a well-accepted marker of PI3K/mTOR pathway activation (Park et al, 2008; Kanaizumi et al 2018; Zhang et al, 2020). Consistent with this, cortical neurons transduced with AAV1-hSYN-PIK3CD exhibited strong pS6 immunostaining in the area of PI3Kδ expression, with clear PI3Kδ and pS6 co-expression (∼38%; Figure 1F, G). Notably, pS6 levels in the transduced hemisphere were significantly elevated compared to the non-injected hemisphere, GFP controls and uninjected controls (Figure 1H-J, F), where pS6 remained at baseline. Together, these results confirm that AAV1-hSYN-PIK3CD drives long-term expression in the cortex and sustains activation of the PI3K/mTOR pathway up to one-year post-injection.

### Long term AAV1-hSYN-PIK3CD expression does not affect glial cell activation

To evaluate the long-term effects of sustained AAV1-hSYN-PIK3CD expression on glial reactivity, we analysed astrocytic and microglial morphology in the motor cortex by immunostaining brain sections for glial fibrillary acidic protein (GFAP), an astroglial lineage marker, and ionised calcium-binding adaptor molecule 1 (Iba1), a microglial marker. Although AAV1 under the hSYN promoter is neuron-selective and does not transduce astrocytes or microglia, it is important to assess whether paracrine signalling or chronic neuronal PI3Kδ overexpression indirectly induces glial activation. We did not observe PI3Kδ expression in either astrocytes or microglia. GFAP immunostaining revealed a modest, non-significant increase in MFI in PI3Kδ-treated animals relative to uninjected controls (Figure 2A-C). This local increase in GFAP signal was spatially restricted to the injection site, and likely reflects mechanical disruption induced astrogliosis rather than PI3Kδ-driven astrocytic activation (Figure S1). Supporting this, GFAP intensity progressively declined with distance from the visible injection site, even in regions where PI3Kδ+ neurons were still present (Figure S1). Similarly, Iba1 immunoreactivity was slightly elevated in PI3Kδ-treated animals compared to uninjected controls (P = 0.1754; Figure 2G-I), although a visible injection site was not detectable with Iba1 staining. We therefore also analysed microglial abundance in the PI3Kδ positive region, of which there were no differences compared to controls (Figure S2). To determine whether these changes were associated with morphological hallmarks of glial activation, we performed Sholl analysis on individual astrocytes and microglia, measuring the number of process-sphere intersections as a function of distance from the soma (Figure 2E, K). Astrocytes in the dense region immediately surrounding the injection site were not analysed as it was difficult to identify individual astrocytes and their respective processes (Figure S1). Both glial cell types exhibited a stereotypical increase in branching with distance from the soma, peaking at ∼15 μm for astrocytes and ∼20 μm for microglia, followed by a decline (Figure 2E, K). Importantly, there were no significant differences in number of intersections at any radius, nor in maximal process length or total intersections per cell between PI3Kδ-treated and control animals (Figure 2F, L). The high number of fine, radially distributed branches is characteristic of a ramified morphology, consistent with a non-activated, homeostatic glial state. These findings suggest that long-term PI3Kδ overexpression in neurons does not elicit chronic astrocyte or microglial activation in the PI3Kδ+ region.

**Figure 2.**
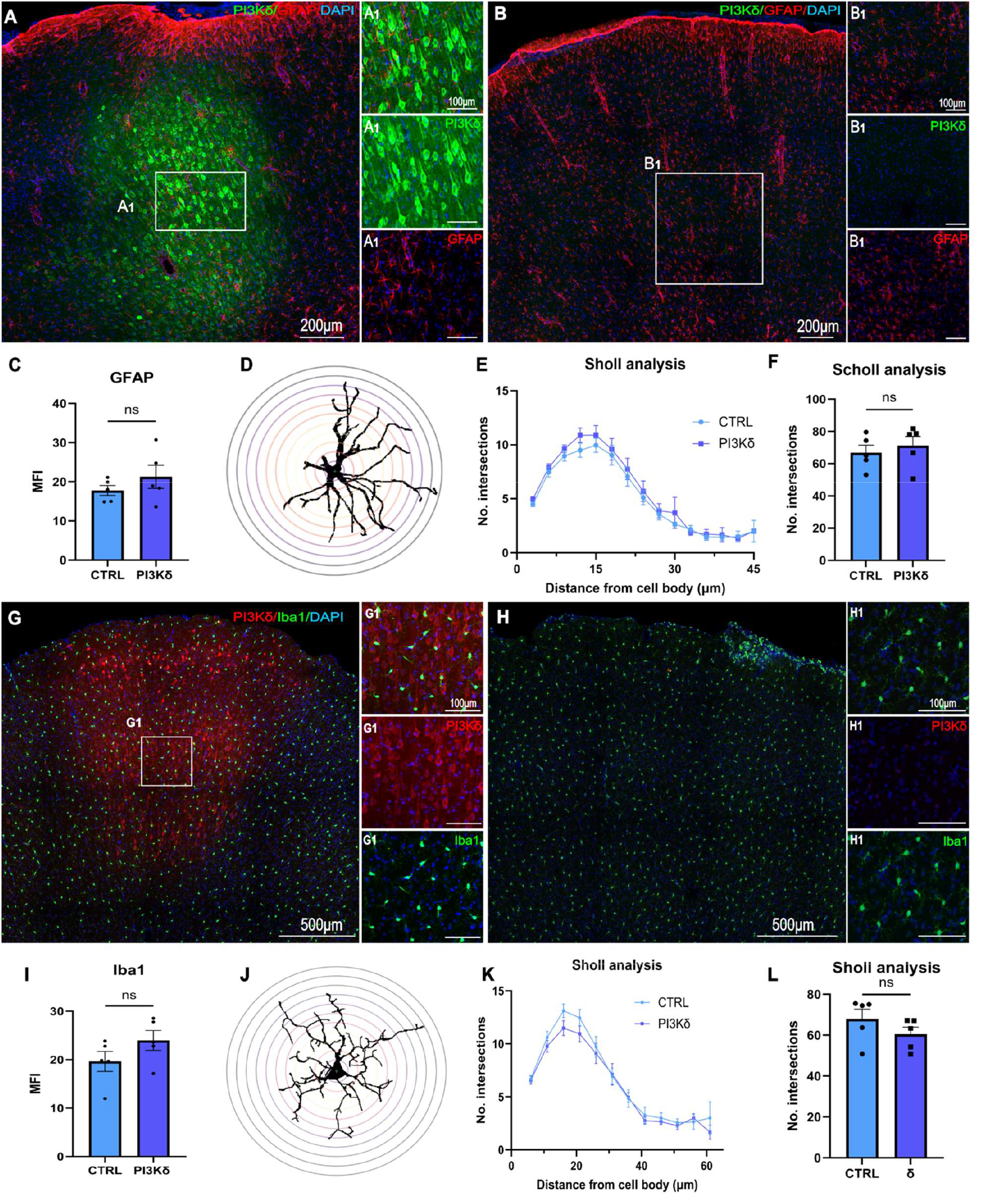
Glial cell mass and morphology remains unchanged following long-term PI3Kδ overexpression. GFAP staining of astrocytes in the motor cortex of AAV1-hSYN-PIK3CD expressing rats (**A**) and controls (**B**). Quantification of MFI of GFAP (**C**) in cortical layer V. Sholl analysis of individual astrocytes, with concentric circles at 3 μm intervals from the soma, colourful dots represent intersections (**D**). Graph of number of intersection points at increasing distances of 3 μm intervals from cell body (**E**) and average number of intersections per cell (**F**). Iba1 staining of microglia in the motor cortex of AAV1-hSYN-PIK3CD expressing rats (**G**) and controls (**H**). Quantification of MFI of Iba1 (**I**) in cortical layer V. Sholl analysis of individual microglia, with concentric circles at 5 μm intervals from the soma (**J**). Graph of number of intersection points at distances moving away from cell body (**K**) and average number of intersections per cell (**L**). Data shown as mean ± SEM, 21 cells/animal were analysed across 3 sections, *n* = 5. Data analysed by T-test or two-way-ANOVA with Tukey’s post hoc test, ns = non-significant.

### No evident PI3Kδ-related brain pathologies or tumours

To further assess the safety of long-term AAV1-hSYN-PIK3CD overexpression, given its established role as an oncogene, we evaluated brain morphology using high resolution ^1^H MRI and histological analysis with Haematoxylin and Eosin (H&E) staining. MRI scans of treated animals revealed no evidence of tumour formation or gross pathological abnormalities. All other structural irregularities observed were confined to regions corresponding to the injection site (Figure 3B1, 2; arrows). H&E staining of coronal brain sections from both PI3Kδ-treated and control animals confirmed preservation of normal cortical architecture. No signs of necrosis, oedema, or cell infiltration were observed. Detailed examination of the superficial cerebral cortex (Figure 3D1, E1) and lateral cerebral ventricle (Figure 3D2, E2) show no differences in cellular histological structure.

**Figure 3.**
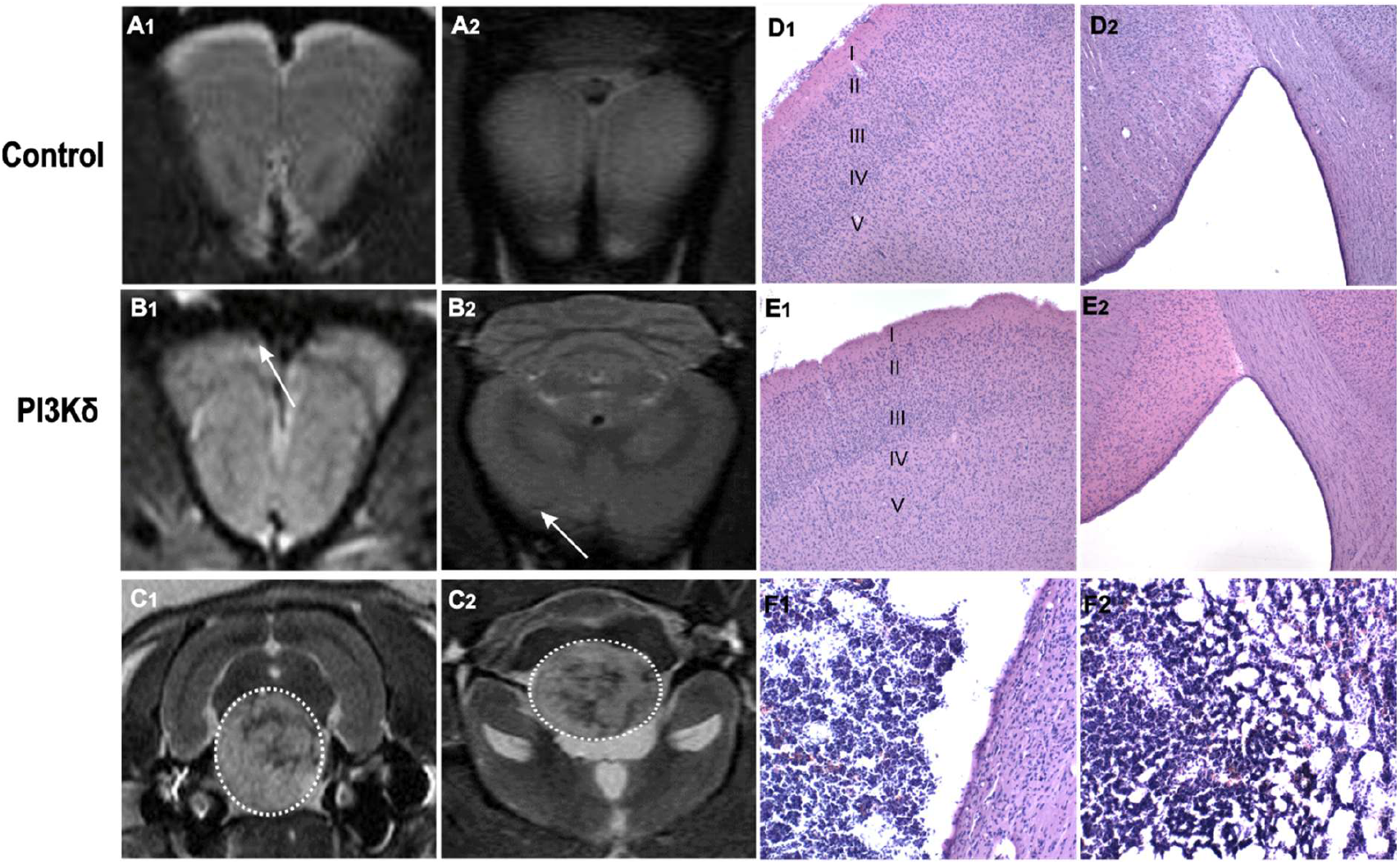
Brain MRI and haematoxylin and eosin staining shows no brain PI3Kδ-related pathologies or tumours. MRI images show absence of tumour formation in both control (**A1**,**2**) and PI3Kδ-treated animals, which sustained minor damage from injection, annotated by white arrows (**B1**,**2**). One tumour was detected in a control animal developing to a volume of 0.21mL, indicated by white dotted lines (**C1**,**2**), H&E staining of coronal brain sections shows normal brain architecture, cortical layers are numbered from superficial (I) to deep (V) (40X magnification) (**D1**,**2, E1**,**2**). Conversely, the detected tumour was compressing adjacent brain tissue (**F1**) and was of abnormal cellular architecture, shown in tumour centre (100X magnification) (**F2**). *n =* 10 per group.

Notably, a tumour with an approximate volume of 0.21 mL was detected in one uninjected control female animal, clearly visible on both axial (Figure 3C1) and coronal (Figure 3C2) MRI scans. The mass appeared to compress the adjacent brain tissue (Figure 3F1) and exhibited abnormal cellular architecture, including signs of reticular reorganisation (Figure F2). No similar abnormalities were found in the remaining control animals or in any PI3Kδ injected animals, suggesting that the tumour was unrelated to PI3Kδ treatment and may instead reflect an age-associated random pathology.

### Sustained PI3Kδ shows no adverse systemic or histopathological effects

In addition to assessing local brain pathology, we evaluated whether long-term PI3Kδ expression exerted any systemic effects by examining overall animal health, including body and organ weight, functional markers and morphology. Data from male and female animals were initially analysed separately and showed no sex-specific differences. Therefore, for conciseness of data presentation, results are presented as combined data for each group. Animal weights were monitored throughout the study, and comparisons between PI3Kδ-treated and uninjected control groups revealed similar trajectories of weight gain over time, with no significant differences observed between groups (Figure 4A). Weight for both groups plateaued at week 40, therefore weighing was concluded at week 45. We also measured the weights of major peripheral organs, of which there were no significant differences between PI3Kδ-treated and uninjected control animals for spleen, lungs, or kidneys (Figure 4A1, 2, 4). Liver weight was significantly heavier in PI3Kδ-treated animals compared to controls (p = 0.048) (Figure 4A3). This may reflect individual variability, metabolic changes, or mild hepatomegaly, although no overt signs of liver pathology, nor any other organ pathology, were detected in gross examination or during tissue collection.

**Figure 4.**
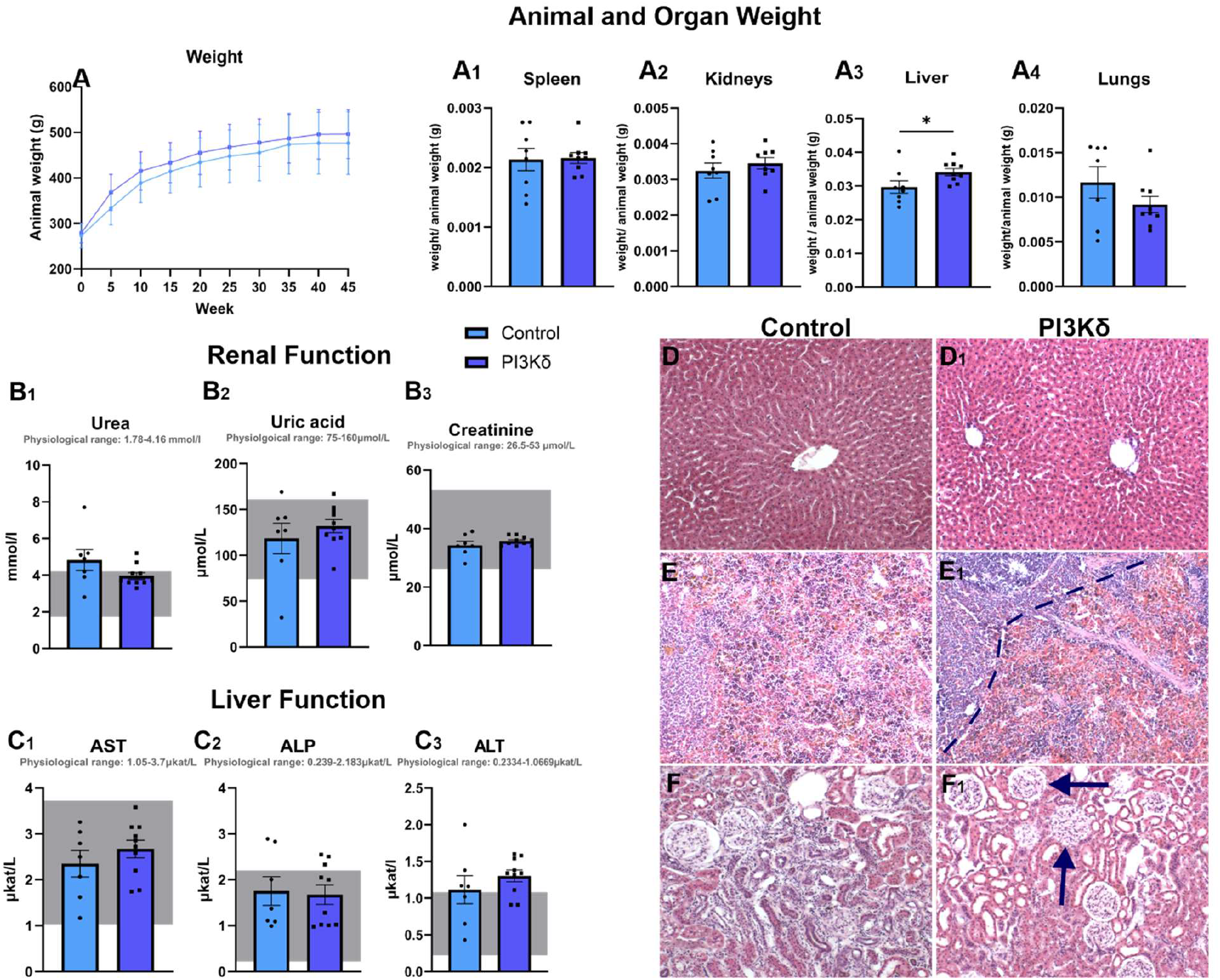
Animal and organ weights, renal and liver function markers, and Haematoxylin and eosin staining of liver, spleen, and kidney following long term AAV1-hSYN-PIK3CD expression. Animal body weight over the duration of 45 weeks (**A**), and organ weights at termination point (**A**_**1**_**-A**_**4**_). Renal function (**B**_**1**_**-B**_**3**_) and liver function (**C**_**1**_**-C**_**3**_) markers taken at 45 weeks. The horizontal grey box indicates nominal physiological range for each parameter. Haematoxylin and eosin staining of controls on left and AAV1-hSYN-PIK3CD treated animals on the right, of liver (**D, D1**), spleen (**E, E1**) with dotted line representing boundary between red and white pulp, and kidney (**F, F1**) with arrows pointing to glomeruli. Data shown as mean ± SEM, ^*^ *p* < 0.05 by T-test, *n* = 7 to 10.

Next, we assessed key biochemical parameters in blood serum, focusing on kidney function (urea, uric acid, creatinine), which together provide an overview of renal filtration efficiency and potential kidney injury (Figure 4B). We also measured liver injury markers: Aspartate Aminotransferase (AST), Alkaline Phosphatase (ALP), Alanine Aminotransferase (ALT), given the increase in liver weight in PI3Kδ-treated animals, and its central role in AAV metabolism and clearance (Figure 4C). Elevations in ALT and AST are commonly used indicators of hepatocellular injury and inflammation, and elevated ALP can indicate cholestatic hepatotoxicity (Thakur et al, 2024). We observed no significant differences between control and PI3Kδ-treated animals for any of the markers, with most serum values falling within the normal physiological range (Figure 4B, C). Minor elevations in urea and ALT levels were noted, as previously reported for ageing rats, but remained close to reference thresholds (Yang et al, 2015). Urinary parameters were also assessed (proteinuria, pH, leukocytes, specific density and glycosuria) and showed no significant differences between uninjected controls and PI3Kδ-treated animals (Figure S3).

To evaluate potential histopathological changes following long-term PI3Kδ treatment, H&E staining was conducted on liver, spleen and kidney tissue sections. Histological analysis of the liver for both control and treated groups revealed normal hepatic architecture, characterised by numerous polygonal hepatocytes arranged in a trabecular pattern, intact sinusoidal architecture and absence of inflammation or fibrosis (Figure 4D, D1). These findings suggest that despite an increase in liver weight and slightly elevated ALT, prolonged PI3Kδ treatment did not induce adverse histopathological effects or compromise liver function. Similarly, H&E staining revealed that both control and PI3Kδ-treated animals exhibited largely normal spleen morphology (Figure 4E). Distinct boundaries were observed between red and white pulp (Figure 4E1). Occasional clusters of disintegrating erythrocytes were present within the red pulp, consistent with splenic erythrocyte turnover. While extramedullary haematopoiesis is typical in foetal and neonatal spleens, small foci can also be observed in adult rat spleens under physiological conditions, and may expand as a compensatory mechanism in response to anaemia, bone marrow dysfunction, or increased haematopoietic demand (Suttie, 2006). However, given the absence of splenomegaly and other signs of erythropoiesis, anaemia, or hematopoietic stress, the observed extramedullary haematopoiesis can be considered a non-pathological, age-related finding in aging Wistar rats (Zhang et al, 2018). Likewise, both control and PI3Kδ-treated animals show a normal histological organisation of the kidney, with several approximately equally sized ovoid glomeruli in contact with collecting ducts of the renal cortex (Figure 4F, F1). Distinct organ related pathologies were observed and include acute tubular necrosis (ATN), hydronephrosis (HN), anaemia (A), and extramedullary haematopoiesis (EMH) and were mainly found in uninjected control rats (Table 1). Only one PI3Kδ-injected animal displayed ATN. All organ related pathologies reported are either a common finding, or associated with ageing in rats (Wang et al, 2014; Hojo et al, 2019).

**Table 1.** Summary of organ related pathologies. Detected pathologies were compared between uninjected control rats and PI3Kδ injected rats, for kidneys, spleen and liver. ATN-acute tubular necrosis, HN – hydronephrosis, A-anaemia, EMH – extramedullary haematopoiesis. No liver pathologies were detected in either PI3Kδ injected or uninjected control animals. No spleen pathologies were detected in PI3Kδ animals, and 3 animals presented ATN of kidney. 2 uninjected control animals presented with spleen and kidney pathologies. *n* = 7-10.

| ORGAN | CONTROL |  |  | PI3Kδ |  |  |
| --- | --- | --- | --- | --- | --- | --- |
|  | HEALTHY (%) | PATHOLOGY (%) | TYPE OF PATHOLOGY | HEALTHY (%) | PATHOLOGY (%) | TYPE OF PATHOLOGY |
| Kidneys | 75 | 25 | 1x ATN, 1X ATN+HN | 70 | 30 | 3X ATN |
| Spleen | 75 | 25 | 1X A, 1X A + EMH | 100 | 0 | - |
| Liver | 100 | 0 | - | 100 | 0 | - |

### Long-term PI3Kδ Expression Does Not Affect Haematological Parameters

Given the broad involvement of PI3Kδ in leukocyte regulation, erythropoiesis, and platelet biology, and its known dysregulation in conditions such as APDS, we next examined whether long-term AAV-mediated PI3Kδ expression produced measurable alterations in peripheral blood cell profiles. The assessed parameters included total leukocyte count and differential white blood cell counts (lymphocytes and relative % of total WBC count, monocytes and relative % of total WBC count, neutrophils, eosinophils, and basophils) (Figure 5A-K). Red blood cell parameters included erythrocytes count, red cell distribution width (RDW), mean corpuscular volume (MCV), reticulocytes, haemoglobin, mean corpuscular haemoglobin (MCH), haematocrit, as well as thrombocytes as a platelet parameter (Figure 5L-S). Data are presented with a horizontal line representing reference ranges for rats 17 weeks of age or older (“Clinical Laboratory Parameters for Crl:Wi (Han) rats”, Charles River Laboratories International, 2008). There were no significant differences between control and PI3Kδ-treated animals for any haematological parameter (p>0.05). RBC parameters typically fell within the reference range for both treated and control animals. However, some WBC parameters, notably lymphocytes relative %, monocytes, and neutrophils (Figure 5C, D, E, F, G), were found to fall outside the reference range. However, this deviation was observed in both control and PI3Kδ-treated animals, suggesting that the alterations are likely attributable to the effects of aging in the rats, which is associated with increased chronic inflammation and immune dysregulation, and an increase in neutrophil to lymphocyte ratio (Ajayi and Olaleye, 2020, Lee et al, 2024).

**Figure 5.**
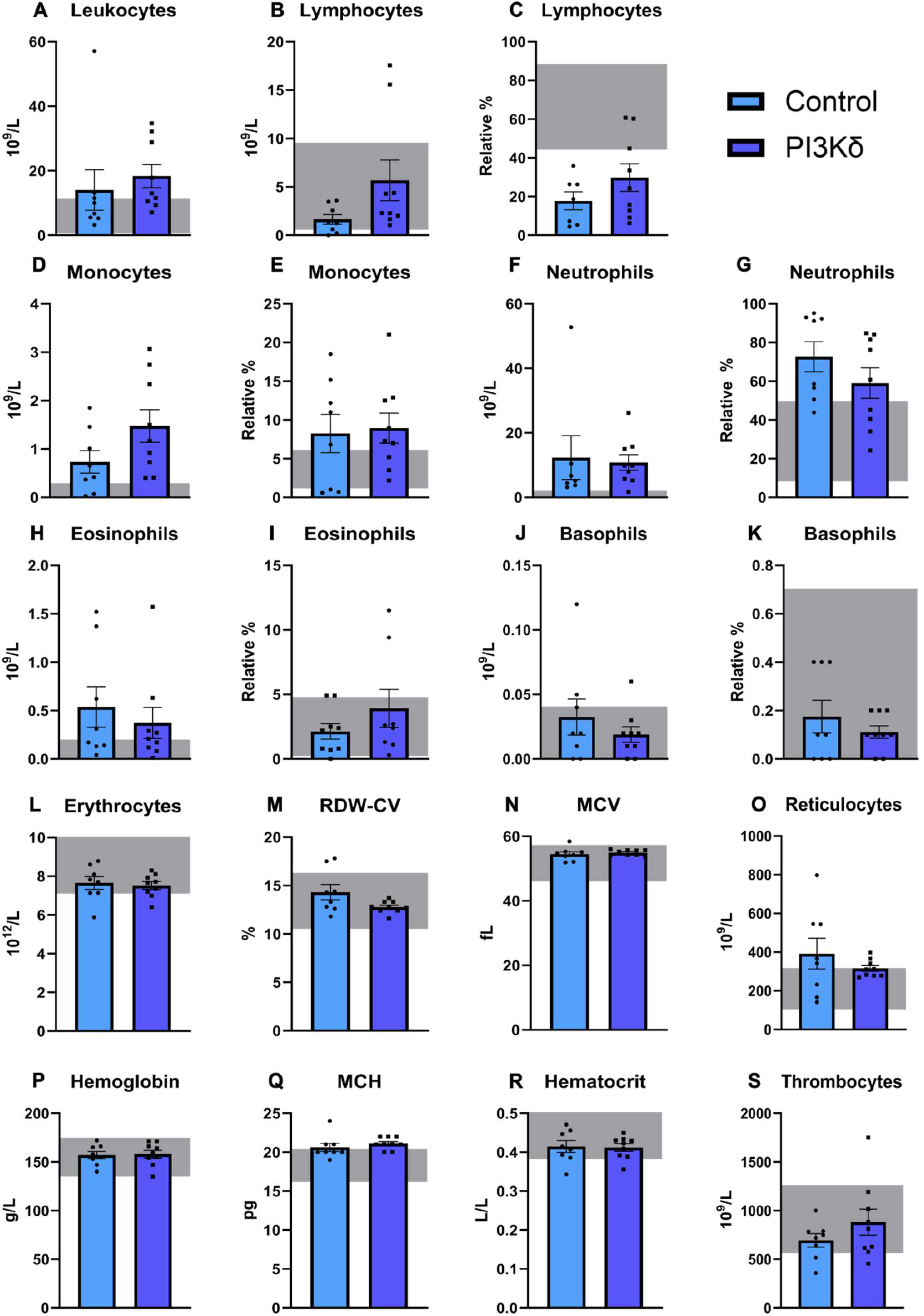
Haematological parameters do not indicate any adverse effects of long term AAV1-hSYN-PIK3CD expression. Assessed parameters: total leukocyte count and differential white blood cell counts (lymphocytes and relative % of total WBC count, monocytes and relative % of total WBC count, neutrophils, eosinophils, and basophils). Red blood cell parameters: red cell distribution width (RDW), mean corpuscular volume (MCV), reticulocytes, haemoglobin, mean corpuscular haemoglobin (MCH), haematocrit, and thrombocytes as a platelet parameter. The horizontal grey box indicates nominal physiological range for each parameter. No significant differences were found between uninjected controls and PI3Kδ-injected animals, by T-test, *n* = 8 to 9.

### Long-term PI3Kδ Expression Does Not Affect Biochemical Parameters

Given the long-term expression of AAV1-hSYN-PI3Kδ in the brain, we evaluated systemic biochemical and metabolic parameters to further assess potential off-target effects and assessment of general safety. Therefore, we measured key blood and serum markers, including electrolytes (calcium, potassium, sodium, phosphorus) that are essential for maintaining physiological functions such as fluid balance, nerve, and muscle activity (Figure 6A-D) (Uzoma et al, 2021). Iron levels were also assessed to exclude iron overload, which can indicate liver injury, or anaemia. In addition, we measured total protein, albumin and serum glucose as indicators of nutritional status, metabolic health, liver and kidney function (Figure 6E-H). As expected, there were no significant differences between control or PI3Kδ-treated animals (p>0.05), as well as the majority of values being within the physiological range. Although reduced serum albumin levels were observed in both groups, this is consistent with previous findings in aging rodents which may reflect age-related hepatic function decline and chronic low-grade inflammation (MacQueen et al, 2011).

**Figure 6.**
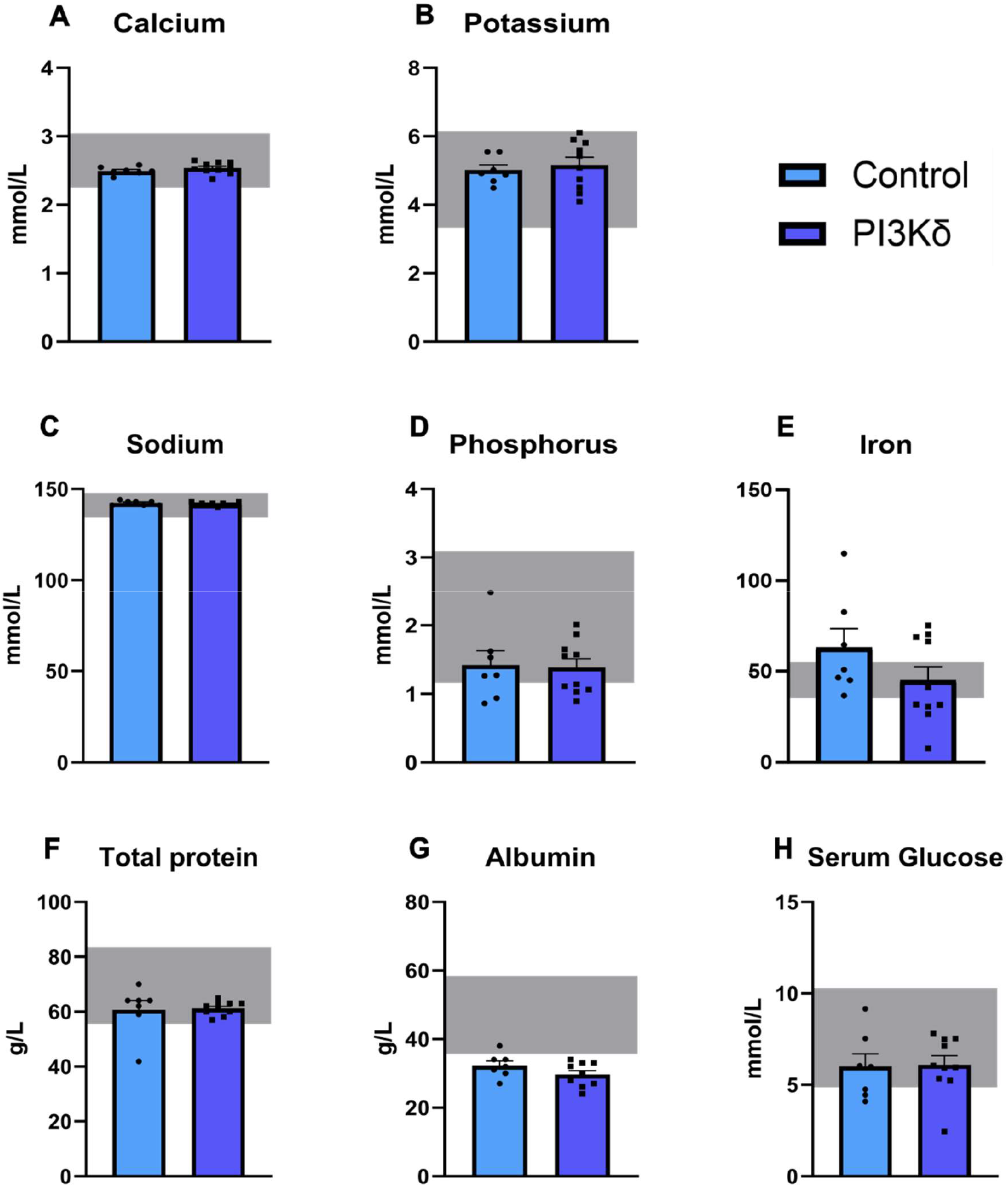
Biochemical parameters do not indicate any adverse effects of long term AAV1-hSYN-PIK3CD expression. Key blood and serum markers, including electrolytes (calcium, potassium, sodium, phosphorus), iron, total protein, albumin and serum glucose were measured (**A-H**). The horizontal grey box represents nominal physiological range for each parameter. No significant differences were found between uninjected controls and PI3Kδ-injected animals, by T-test, *n* = 7 to 10.

## Discussion

Although there remains no cure for spinal cord injury, advances in many gene therapies hold a promising future for patients. One such advancement surrounds the PI3K pathway. In this study, we aim to evaluate the long-term consequences of sustained PI3Kδ overexpression in the adult rat brain following AAV1-hSYN-PIK3CD intracortical delivery.

Previous studies have raised concerns regarding the potential ‘waning’ of AAV-mediated transgene expression over time, attributed to mechanisms such as degradation of episomal DNA, epigenetic silencing via transgene or promoter methylation, and progressive neuronal loss due to aging or neurodegeneration (Prösch *et al*, 1996, Das et al, 2022). For instance, Jackson et al. (2016) reported a marked attenuation of hSYN-driven GFP expression at 22 weeks compared to 4 weeks following intravenous administration to neonatal rats. By contrast, Klein et al. (2002) demonstrated that intracranial delivery of AAV2-CAG-GFP to rats resulted in stable neuronal transgene expression for up to 18 months. However, there remains a lack of studies specifically evaluating the long term (> 6 months) expression following intracranial delivery of AAVs, specifically AAV1, under the synapsin promoter in rats.

For applications requiring direct targeting of cortical neurons, stereotaxic cortical injection offers distinct advantages. This approach enables accurate targeting of the desired cortical area while achieving therapeutic transgene levels with low viral doses, reducing the immune responses, peripheral tissue distribution, and avoiding the issue of pre-existing neutralising antibodies associated with systemic delivery (Huda et al, 2014; Chowdhury et al, 2024; Jeune et al, 2013). It is also important to note that many biodistribution and expression studies often rely on fluorescent reporters, such as GFP (Hutson et al, 2011, Yáñez-Muñoz, 2006, Hollidge et al, 2022). While useful for mapping transduction patterns, it is not completely reliable since AAVs carrying different transgenes have different expression kinetics, protein stability and biological effect (Suarez-Amaran et al, 2025, Choi et al, 2014). Considering the highly invasive nature of intracortical delivery, repeated dosing is impractical, making it best suited for single-dose administration. Confirming stable long-term transgene expression with hSYN-AAV1 is essential for translational CNS gene therapy development. Therefore, in this study we first confirmed that AAV1-hSYN-PIK3CD expression persisted for one-year post-injection, with expression intensity comparable to that reported previously at 16-weeks (Karova et al, 2025). The turnover rate of a target cell can largely influence the durability of AAV-mediated gene expression, therefore since neurons are post mitotic and the AAV genome persists predominantly as episomal circular concatemers, the lack of dilution by mitosis makes this sustained MFI not surprising (Muhuri et al, 2022).

To accurately assess effects of sustained neuronal PI3Kδ, it is essential to confirm whether long term AAV1-hSYN-PI3Kδ driven overexpression resulted in persistent activation of the downstream PI3K signalling pathway. We know this pathway is regulated with several feedback mechanisms and a multi layered network of molecular brakes and amplifiers (Chandarlapaty et al, 2012; Fruman et al, 2018), however we do not know if or how these molecular feedbacks would respond to sustained overexpression of PI3Kδ, especially in neurons, where continued activation of this pathway is unusual and much less studied (Wright et al, 2024). One challenge in the development of inhibitors that target the PI3K/Akt/mTOR pathway is the emergence of drug resistance caused by unintended pathway reactivation through intrinsic and extrinsic feedback loops (Wright et al, 2021). For instance, activation of mTORC1 and S6K leads to phosphorylation of the insulin receptor substrate (IRS), which suppresses IR receptor expression and signalling. This phosphorylation inhibits IRS-1 function, creating a negative feedback loop that limits further PI3K/Akt activation (Tanti and Jager, 2009). Preclinical cancer models and clinical trials have shown that inhibition of mTORC1/S6K using rapamycin analogs results in increased in Akt phosphorylation by PDK1 and MTORC2. This effect arises from relief of S6K-mediated negative feedback on IRS-1, leading to enhanced PI3K signalling and compensatory Akt activation, which may limit the therapeutic efficacy of MTORC1 inhibition alone (O’Reilly et al, 2006). Importantly, it has also been shown that in both physiological and oncogenic settings, activation of the PI3K/Akt/mTOR pathway drives translation of the PTEN protein, serving as a compensatory mechanism to maintain homeostatic control of the pathway (Mukherjee et al, 2022). In the context of neurobiology, increased pS6 immunostaining, the substrate of S6K and well accepted marker of PI3K/mTORC1 pathway activation, has been observed in mice with PTEN deletion, showing qualitatively similar levels at both 3 and 15 months of age (Gallent and Steward, 2018). Here we show that pS6 is still upregulated in neurons expressing PI3Kδ in the injected cortex, with PI3Kδ^+^pS6^+^ cells only slightly less than that reported after 16 weeks (Karova et al, 2025). This confirms that sustained expression of PI3Kδ in cortical neurons maintains activation of the PI3K/Akt/mTOR pathway.

It has previously been shown that overactivation of the PI3K pathway, through viral expression of Akt3 or PTEN deletion, results in neuronal soma hypertrophy, the former showing pS6^+^ cell diameter nearly double that of controls, and the latter showing a continued increase in cell size between 12 and 15 months after treatment (Campion et al, 2022; Gallent and Steward, 2018). Notably, high-titre Akt3 overexpression also induces spontaneous seizure activity and neuronal hypertrophy across multiple neuronal subtypes, a well-established pathological feature associated with spontaneous epileptic seizures, linking excessive PI3K-mTOR activation to aberrant network excitability (Lasarge and Danzer, 2014). Here, we report that sustained overexpression of PI3Kδ increased soma size compared to GFP controls, likely due to pS6 induced cellular hypertrophy, however only by an average of 1μm, with neuron soma size still falling into reported range for adult rats. We therefore did not observe the drastic increase reported following Akt3 expression or PTEN deletion, neither did our animals present seizures. Our findings highlight an important distinction between PI3Kδ overexpression and other manipulations of the PI3K/Akt/mTOR pathway. Although it is not entirely known why, these differences are likely due to the bypassing or removal of some of the homeostatic brakes and feedback systems. Unlike PTEN deletion or Akt3 overexpression, which can activate the pathway independently, PI3Kδ overexpression still relies on upstream RTK receptor activation to exert downstream effects, preserving the endogenous regulatory control.

Arguably, one of the most important analyses in this study were the MRI and histological assessments of brains to determine whether long term PI3Kδ overexpression leads to tumour formation. In this study, we report complete absence of tumours in PI3Kδ expressing animals, while one tumour was detected in a control female animal. This is in line with previous findings that although at very low incidence, spontaneous brain tumours do occur in aged Wistar rats (Bertrand et al, 2014).

Tumours arising from postmitotic cells like neurons may develop through mechanisms of dedifferentiation and deregulation of signalling, notably of the oncogenic MAPK pathway (Bahar et al 2023; Pekmezci et al, 2018; Gorodezki et al, 2024). More commonly, these tumours have a mixed neuronal and glial origin (glioneuronal), with the latter dividing cell type more prone to transformation. These tumours present as low grade, seldom clinically aggressive resulting mainly in symptoms of epilepsy, seizures and headaches in response to built-up pressure on the surrounding nervous tissue (Martinoni et al, 2023). Neuronal or glioneuronal tumours are rare, accounting for approximately 1% of all primary CNS tumours while glioblastoma accounts for 32% of primary brain tumours (Vaz et al, 2022; Nunno et al, 2024; Thakkar et al, 2014). Despite low incidence of neuronal or glioneuronal tumours, PI3Kδ overexpression poses legitimate safety concerns regarding its long-term effect on neurons or risk of off target glial expression and subsequent consequences, given that aberrant activation of the PI3K pathway represents one of the most common molecular events in human cancers (He et al, 2021; Shin et al, 2002). In this study, we confirm that expressing PI3Kδ under the hSYN promoter resulted in strictly neuron-selective expression, mitigating off target glial expression where PI3Kδ could potentially drive uncontrolled proliferation. Since mature neurons are post-mitotic and lack the ability to re-enter the cell cycle, elevated PI3K signalling cannot induce neuronal proliferation (Nieuwenhuis et a*l*, 2020). Although there has previously been some concern regarding cancer as a result of long term AAV gene transfer, the integration frequency of AAVs remains rare (0.1-1%) and so the risk of insertional mutagenesis is very low (Donsante et al, 2001; Hanlon et al, 2019). To date, there is no evidence of tumour formation in the CNS due to AAV1-hSYN mediated gene delivery (Hanlon et al, 2019). Results of this study confirm, that long term expression of PI3Kδ oncogene was observed only in neurons and did not cause development of brain tumours despite sustained signalling.

Another critical consideration for recombinant AAV (rAAV) - mediated gene transfer in the brain is the immune response triggered by either the vector itself, or by transgene expression. Therefore, we characterised the local inflammatory response by analysing astrocytes and microglia. We found that there was an increase in MFI for both glial types when comparing the injected hemisphere with uninjected controls, around the layer V cortical neurons for both astrocytes and microglia. For astrocytes this increase can clearly be attributed to the area immediately surrounding the injection site, as astrocytic density decreases with distance from the needle track, in line with previous studies (Hutson et al, 2011; Reimsnider et al, 2007). A clear injection site was not visible in microglial staining, but cell densities showed no differences between groups. Sholl analysis of both glial cell types in the PI3Kδ transduced area confirmed that this increase in MFI was not associated with activation of glial cells and is likely the response to intracortical injections rather than the presence of AAV1 or neurons overexpressing PI3Kδ.

To evaluate potential systemic and physiological consequences of long term AAV1-hSYN-PI3Kδ expression, we assessed a comprehensive panel of haematological and biochemical parameters, including those indicative of organ health. The liver is a particular focus in AAV safety assessments, as it serves as the primary site of vector clearance and immune activation. In this study, we detected a modest increase in liver weight in PI3Kδ-treated animals. However, H&E staining and measurements of liver function showed no evidence of histopathological abnormalities or impaired hepatic function. We did detect a slight elevation of ALT above the physiological range, which generally indicates hepatocyte injury or stress (Vazquez et al, 2020). However, since this elevation was detected in both PI3Kδ and uninjected control animals, the underlying cause can more likely be attributed to ageing associated decline in liver function (Barcena et al, 2021). A slight increase in liver weight compared to uninjected controls may reflect mild hepatocellular hypertrophy due to potential vector uptake and clearance from venous drainage or cerebrospinal fluid outflow, without causing a pathological change. Notably, AAV1 demonstrates a greater neuronal tropism and reduced hepatic transduction compared to AAV9 (Weber-Adrian et al, 2021). When combined with the targeted intracortical delivery and use of the neuronal selective promoter hSYN, which would restrict transgene expression outside of neurons, significant off-target effects are unlikely. We specifically assessed a wide panel of haematological parameters, such as leukocytes and erythrocytes, given PI3Kδ central role in immune regulation, as well as key biochemical parameters to confirm no unexpected physiological effects. Importantly, we found no signs of adverse off target effects with any other haematological or biochemical parameter, nor with H&E staining. These results are in agreement with a previous study that assessed a similar comprehensive panel of haematological and biochemical parameters, animal weights, micro and macroscopic analysis of organs, following long term AAV2 delivery to rat striatum (Hackett et al, 2005).

Taken together, our findings demonstrate that long-term AAV1-hSYN-PI3Kδ expression in the adult rat cortex is well tolerated, with stable neuronal transgene expression and sustained pathway activation in the absence of adverse cellular, systemic, or oncogenic effects. These results support the safety and durability of neuron-specific PI3Kδ overexpression using AAV1, providing a strong foundation for further preclinical evaluation of targeted gene therapies in spinal cord injury and related neurological disorders.

## Acknowledgements

This research was supported by the **International Foundation for Research in Paraplegia (IRP) P186, MEYS CR INTER-ACTION LUAUS25_LUAUS25141, OPJAK EXREGMED CZ.02.01.01/00/22_008/0004562**, and by the Microscopy Service Centre of the Institute of Experimental Medicine CAS supported by the **MEYS CR** (**LM2023050 Czech-Bioimaging**), which includes support by the Ministry of Education, Youth and Sports of the Czech Republic (**Research Infrastructure NanoEnviCZ, LM2018124**) and by The European Union – European Structural and Investment Funds in the frame of the Research Development and Education – project Pro-NanoEnviCZ operational program (**Project No. CZ.02.1.01/0.0/0.0/16_013/0001821**). For Open Access, a CC BY 4.0 public copyright license is applied to any Author Accepted Manuscript (AAM) version arising from this submission.

## Notes

### Competing Interest Statement

The authors have declared no competing interest.

