## Supplemental files (S1-S3) for "Long-Term Expression and Safety of AAV1-Mediated PI3Kδ Overexpression in the Adult Rat Cortex"

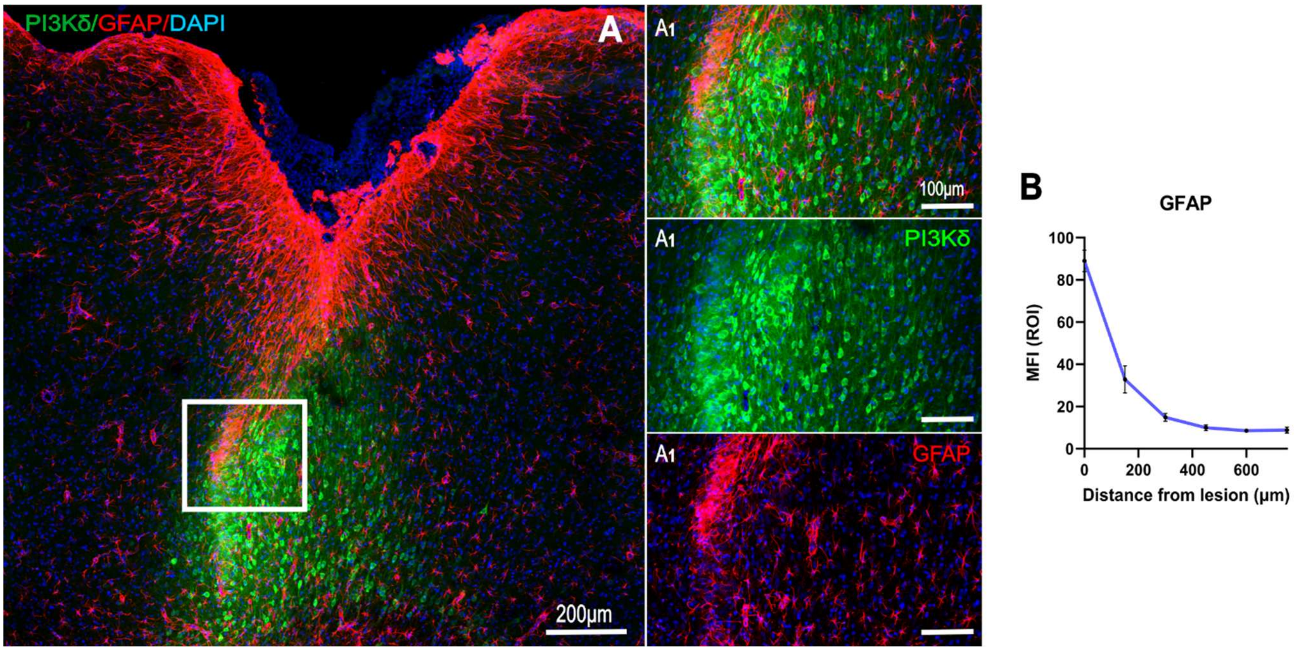


**Figure S1. MFI of astrocytes, related to figure 2.** GFAP staining of astrocytes in the motor cortex of AAV1-hSYN-PIK3CD injected rats (**A**). MFI of GFAP with increasing distance of 200 µm from the visible injection site (**B**). Data shown as mean ± SEM, 3 images were analysed per animal, *n* = 5

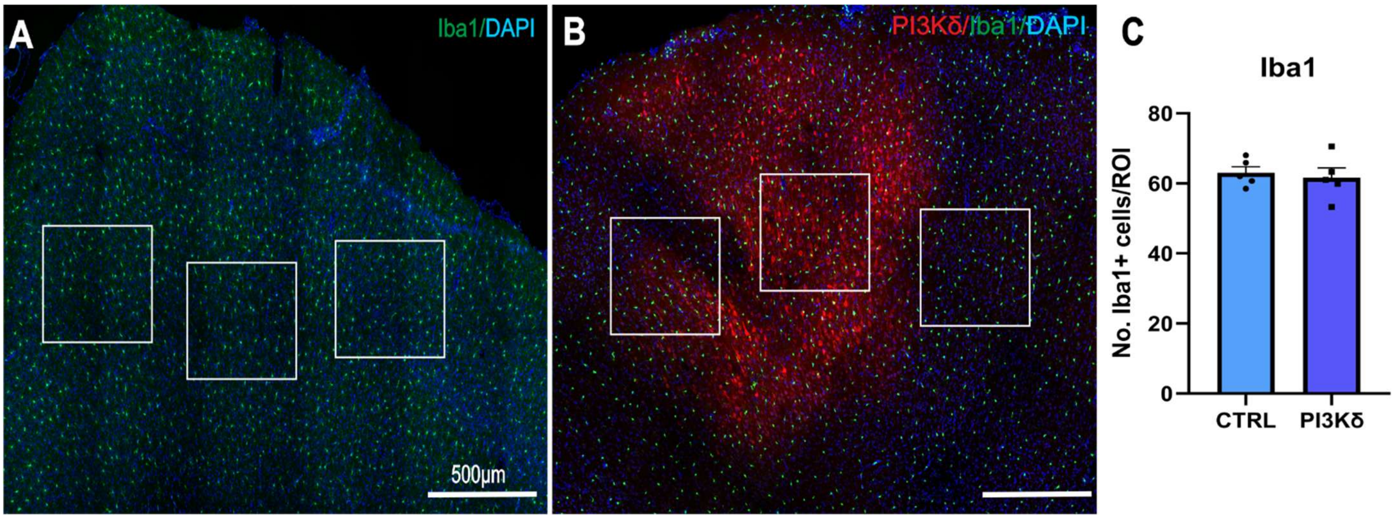


**Figure S2. Iba1 cell density, related to figure 2.** Iba1 staining of microglia in the motor cortex of uninjected controls (**A**) or AAV1-hSYN-PIK3CD injected rats (**B**). White squares represent ROIs 500 µm x 500 µm from which microglial numbers were counted and averaged across 3 ROIs per image. Data shown as mean ± SEM, 3 images were analysed per animal, *n* = 5.

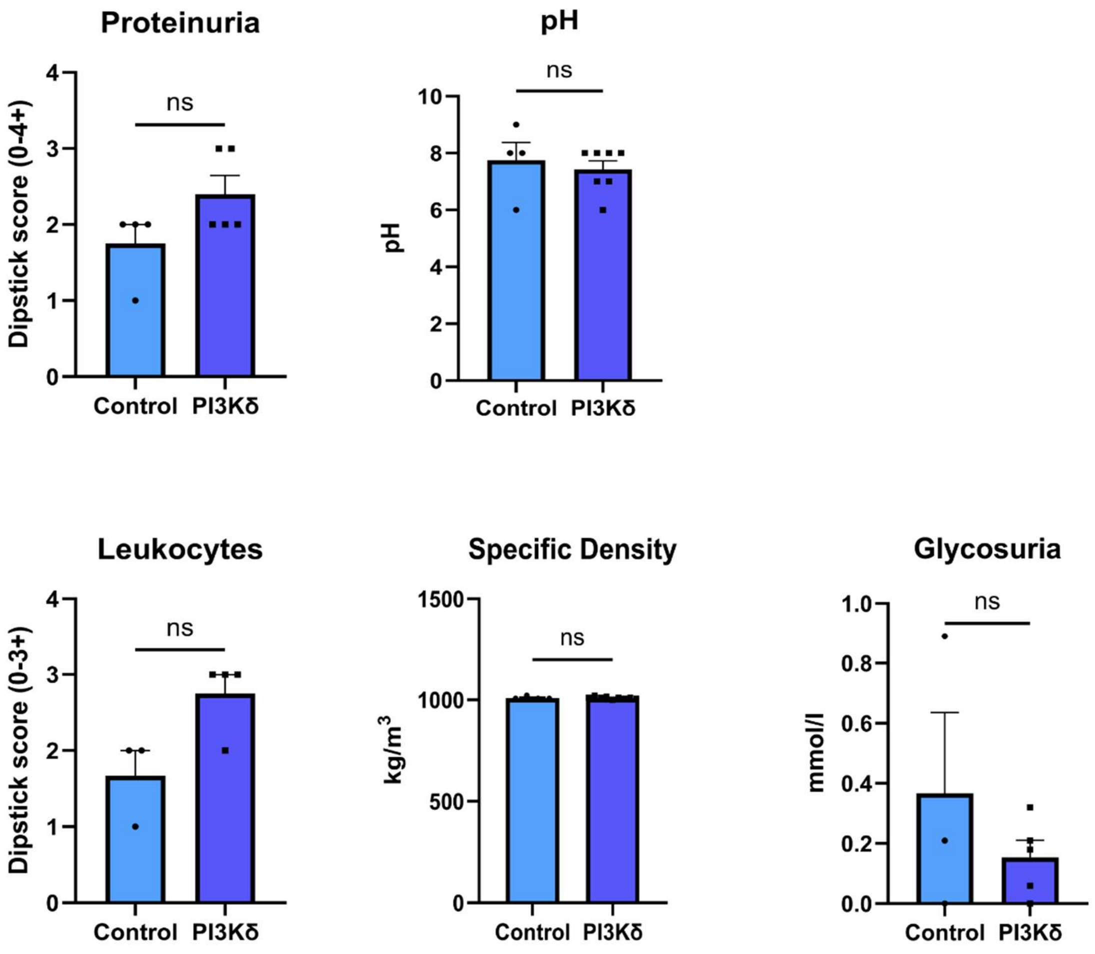


**Figure S3. Urinary parameters do not indicate any adverse effects of long term PI3Kδ expression.** Assessed parameters: proteinuria, pH, leukocytes, specific density and glycosuria measured in animal urine 1 year after AAV1-hSYN-PIK3CD injections. No significant differences were found between uninjected controls and PI3Kδ-injected animals, by T-test, *n* = 3 to 7.
